# Priming the pump: Enhanced nitrite release in response to a nitrate pulse by nitrogen-limited *Prochlorococcus*

**DOI:** 10.1101/2025.05.13.653865

**Authors:** Paul M. Berube, Trent LeMaster, Sallie W. Chisholm

## Abstract

*Prochlorococcus* is a diverse and widespread cyanobacterium with significant contributions to the marine nitrogen and carbon cycles. Some *Prochlorococcus* reduce and divert up to 20-30% of the nitrate that they take up to external pools of nitrite. Given that nitrite is a central intermediate of the nitrogen cycle and *Prochlorococcus* is highly abundant in nitrogen-limited waters, our goal was to advance our understanding of nitrite cycling in the context of nitrogen limitation. Here we observe that nitrate-limited *Prochlorococcus* have cell-specific nitrite production rates that are approximately a magnitude higher than nitrogen-replete *Prochlorococcus* when challenged with a pulse of nitrate. Nitrite production rates are unchanged or depressed during light and cold shocks, suggesting that nitrate is not used as an alternative electron acceptor to mitigate the impacts of excess photochemically generated electrons. These results suggest that in regions where phytoplankton growth is limited by nitrogen, *Prochlorococcus* cells could be primed to transform substantial quantities of nitrate into extracellular pools of nitrite during episodic upwellings of nitrate-rich water. Given that nitrite is an important intermediate in the nitrogen cycle, these results have ramifications for our understanding of nitrogen cycling in nitrogen-limited open ocean ecosystems.

## Introduction

*Prochlorococcus* is the numerically dominant photosynthetic organism in the vast subtropical gyres of the world’s oceans. Together with its close relative, *Synechococcus*, it is responsible for a sizable fraction of marine carbon fixation (Flombaum et al., 2013). These ocean gyres are oligotrophic — i.e., they are low in nutrients (particularly N, P, and Fe) and have low primary productivity. Subtropical gyres are among the largest biomes by area on the planet and thus have a significant carbon fixation footprint. *Prochlorococcus* thrives in these environments, in part due to the number of adaptations it has acquired since its evolutionary radiation from *Synechococcus*. For instance, *Prochlorococcus* has an open pangenome with many accessory genes spread across the full extent of its diversity (Biller et al., 2015). Its genomes are thus optimized to distinct habitats, such as the phosphorus-limited North Atlantic Subtropical Gyre (Coleman & Chisholm, 2010; Martiny et al., 2006). The *Prochlorococcus* that are selected for in different ocean regimes have complements of accessory genes that are so finely tuned to environmental pressures that these genes can be used to predict the proximal limiting nutrient where *Prochlorococcus* are abundant (Garcia et al., 2020; Ustick et al., 2021). It has further been hypothesized that more recently emerged lineages have evolved photosynthetic electron transport chains that maximize electron flow, which in turn facilitates nutrient uptake at low concentrations (Braakman et al., 2017).

Large swaths of the oceans’ subtropical gyres, where *Prochlorococcus* is the dominant primary producer, are characterized by concentrations of inorganic nitrogen that are low enough to limit primary productivity (C. M. Moore et al., 2013). In addition to using ammonium as a nitrogen source, many clades of *Prochlorococcus* possess accessory genes encoding assimilation pathways for nitrite, nitrate, urea, cyanate, and amino acids (Berube et al., 2015; Kamennaya & Post, 2011; Kamennaya et al., 2008; L. R. Moore et al., 2002; Yelton et al., 2016; Zubkov et al., 2004). Within the LLI clade of *Prochlorococcus* — the most abundant low-light adapted clade — there are 3 known functional types with respect to nitrate and nitrite metabolism (Berube et al., 2019). Some cells have lost the ability to use nitrate, but can still assimilate the more reduced nitrite into biomass. Others can use both nitrate and nitrite. Among the LLI *Prochlorococcus* that can use both, one functional type has retained a nitrite-specific transporter (FocA/NirC), while another functional type has lost the gene encoding the nitrite-specific transporter and further encodes a divergent copy of the nitrite reductase, NirA (Berube et al., 2019).

A representative strain that lacks the nitrite-specific transporter and encodes the divergent (type II) nitrite reductase has been shown to release sizable quantities of nitrite during exponential growth on nitrate in batch laboratory cultures (Berube et al., 2023). Approximately 20-30% of nitrogen transported into the cell as nitrate was reduced and released back to the extracellular environment as nitrite. This is a relatively large loss of N for an organism adapted to the nutrient-poor subtropical ocean. This phenomenon occurs at low concentrations of nitrate similar to those observed in the sunlit upper water column of the North Pacific Subtropical Gyre during intermittent upwelling events that inject nitrate from deeper waters into the surface waters (Berube et al., 2023; Johnson et al., 2010). We hypothesize that this functional type of LLI *Prochlorococcus* may be adapted to persistent low nitrate availability — i.e., the transporters and reductases of its nitrate and nitrite assimilation pathway may have evolved high affinity for their substrates but low throughput or transport/reaction velocity. Under conditions of a nitrate pulse that might exceed the Ks (half-saturation constant) for growth on nitrate, this pathway would likely experience a bottleneck that results in an accumulation of nitrite in the cell that is subsequently released. Here we evaluate this hypothesis alongside an alternative hypothesis that nitrate could be reduced to nitrite by excess photochemically generated electrons as a photoprotective mechanism during light or temperature shock.

## Results and Discussion

### The potential for incomplete assimilatory nitrate reduction is magnified in nitrogen-limited

***Prochlorococcus***. The availability of nitrogen often limits the growth of *Prochlorococcus* in the wild. In the context of this defining feature of *Prochlorococcus*’ evolution, we hypothesized that the MIT0917 strain releases high concentrations of nitrite during nitrate-replete growth because it carries a NirA nitrite reductase enzyme optimized to nitrogen-limited conditions (Berube et al., 2023). In this scenario, *Prochlorococcus* MIT0917 could be prone to a bottleneck in the nitrate assimilation pathway due to the kinetics of NirA whereby the catalytic throughput of the enzyme would be less than that of the NarB nitrate reductase, with excess nitrite being released from the cell. If so, nitrogen-limited growth — with expected high expression of nitrate assimilation proteins (primarily the NapA nitrate/nitrite transporter and reductases) — could prime the cells for elevated nitrite release when challenged with nitrate. To assess this, we grew the MIT0915 and MIT0917 strains in nitrate-limited chemostats (Figures 1, Supplementary Figure S1) at two dilution rates to assess the response of cultures to variable degrees of nitrogen-limitation. Following a switch in dilution rate — and hence the nitrogen-limited growth rate — from 0.25 d^-1^ to 0.5 d^-1^, cell numbers declined as expected given the higher cellular N quota required for elevated growth rate (Droop, 1968, 1973, 1974). Nitrate concentrations in all chemostats were undetectable during steady state at the final target dilution rate of 0.5 d^-1^.

**Figure 1.**
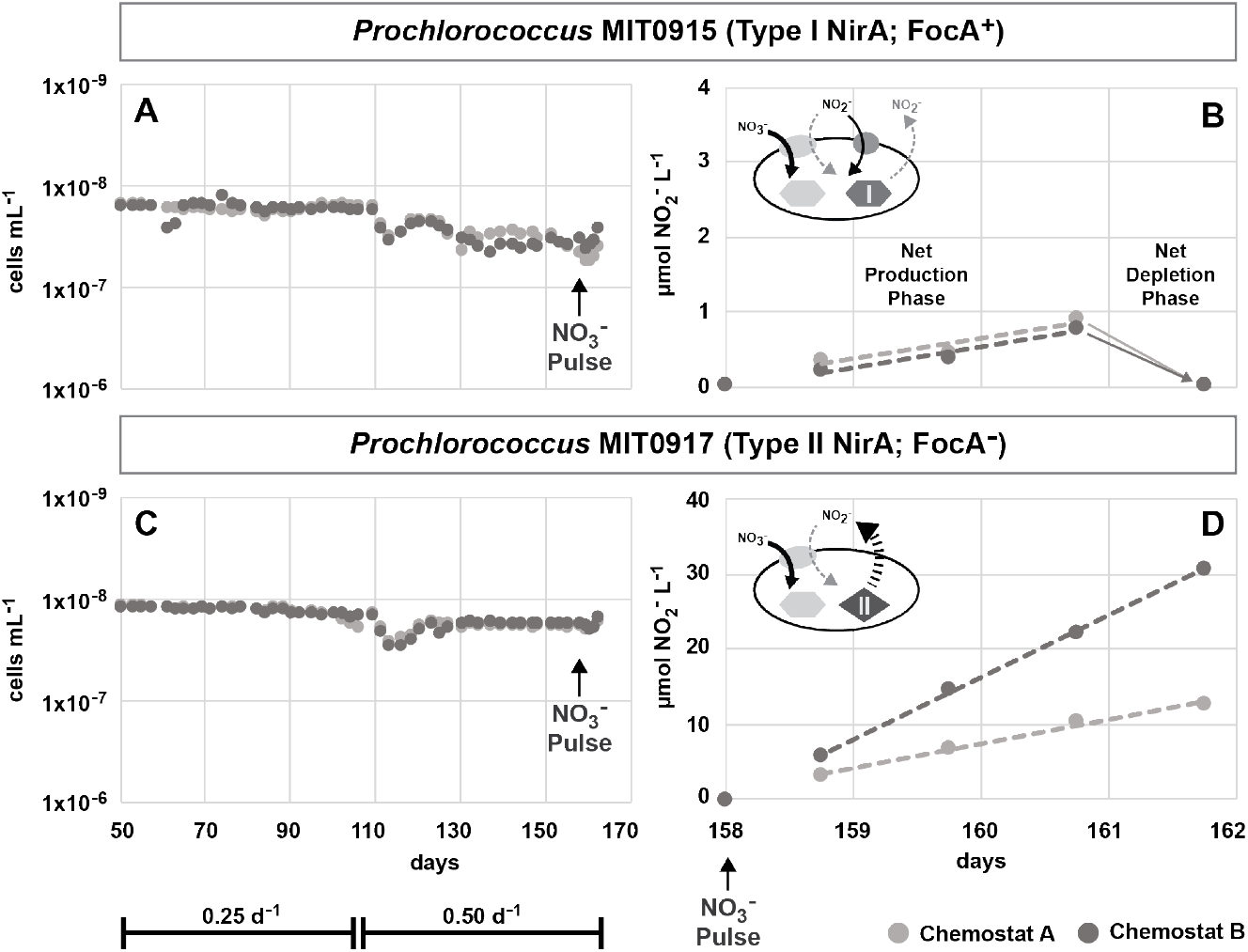
*Prochlorococcus* cell concentrations for MIT0915 and MIT0917 during continuous culture in nitrate-limited chemostats and changes in extracellular nitrite in response to a nitrate pulse. *Prochlorococcus* MIT0915 (A, B) and *Prochlorococcus* MIT0917 (C, D) were grown in chemostats with nitrate as the limiting nutrient in artificial seawater. Cell concentrations (A, C) were profiled in duplicate chemostats for each strain at dilution rates of 0.25 d^-1^ and 0.5 d^-1^. On day 158, the chemostats were spiked with nitrate, after which there was a modest increase of cell concentrations over the subsequent 4 days (A, C); 12-13% increase for MIT0917 and 18-27% increase for MIT0915. Nitrite concentrations in the chemostats for MIT0915 (B) and MIT0917 (D) are shown for the final 5 days, with dashed lines indicating the data points used to determine the effective nitrite production rate in each chemostat.

To assess cell-specific nitrite production rates in nitrate-limited cells at a growth rate of 0.5 d^-1^, we spiked each chemostat with 800 μM of sodium nitrate and measured the concentration of nitrite over a 4 day time course (Figure 1). The response of nitrate-limited MIT0917 cells to a nitrate pulse was particularly notable; the cell-specific nitrite production rates were 1.53 × 10^−7^ (± 0.09 × 10^−7^) nmol nitrite cell^-1^ d^-1^ in one of the duplicate chemostats containing MIT0917 and 3.60 × 10^−7^ (± 0.50 × 10^−7^) nmol nitrite cell^-1^ d^-1^ in the other chemostat. These rates are up to 6-fold higher than those previously observed (Berube et al., 2023).

We also observed detectable nitrite production by MIT0915 for the first time (Figure 1), with the caveat that heterotrophic bacteria were observed in the MIT0915 chemostats (Supplementary Information). Previously, incomplete nitrate assimilation with concomitant nitrite release was only observed in the MIT0917 strain with the Type II nitrite reductase (Berube et al., 2023). While the nitrite concentrations in the chemostats containing MIT0915 remained low (*<*1 μmol L^-1^ nitrite) and ultimately declined to *<*0.05 μmol L^-1^ nitrite during the final 24 hours of the experiment — possibly due to uptake of nitrite via the nitrite-specific FocA transporter found in MIT0915 or by one of the heterotrophs present in the cultures — these data suggest that incomplete assimilatory nitrate reduction is found in LLI *Prochlorococcus* with a Type I NirA nitrite reductase. Assembly of the dominant *Aurantimonas* strain (Supplementary Materials and Methods) indicated that it encoded the machinery for the uptake and reduction of both nitrate and nitrite. Thus, this heterotrophic partner may have been a sink for any nitrite produced by MIT0915. If so, the apparent rates of nitrite accumulation in nitrate-limited cultures of MIT0915 following a nitrate pulse may be an underestimate of the actual nitrite release rates. Regardless, this does not change the conclusion that nitrogen-limited growth increases nitrite production potential due to incomplete assimilatory nitrate reduction in the MIT0917 strain and possibly in the MIT0915 strain as well.

### Light and cold shocks depress nitrite production by *Prochlorococcus*

While it is clear that the potential for nitrite production in response to a nitrate pulse is greatly elevated in cells that are in a nitrogen-limited state, we also hypothesized that nitrate reduction could function as a photochemical safety valve. Nitrate is known to be used as an electron acceptor to dissipate excess energy captured by photosystems in marine phytoplankton (Glibert et al., 2013; Lomas & Glibert, 1999, 2000). Excess energy can accumulate as a result of an increase in irradiance or a decrease in temperature — i.e., because of limited photosynthetic capacity or temperature-dependent constraints in the metabolic pathways receiving photochemically generated electrons. The reduction of molecular oxygen and subsequent generation of reactive oxygen species (ROS) — which inactivate photosystems — can result from elevated cellular electron pressure (Shimakawa & Miyake, 2018).

*Prochlorococcus* is particularly vulnerable to ROS because it lacks the catalase enzyme which detoxifies hydrogen peroxide (J. J. Morris et al., 2008, 2011, 2012). Many *Prochlorococcus* cells, including those belonging to the LLI clade, possess a plastid terminal oxidase (PTOX) which uses molecular oxygen as a terminal electron acceptor, thus serving as a primary safety valve (Bagby & Chisholm, 2015). Recent theory also postulates that *Prochlorococcus* have evolved to maximize electron flux through its photosynthetic apparatus to facilitate nutrient acquisition at ever lower concentrations (Braakman et al., 2017). Thus, we hypothesized that nitrate may serve as an alternative electron acceptor during periods of elevated energy pressure (e.g., supplementing the existing safety valves that use either molecular oxygen or carbon dioxide as electron acceptors).

To assess the potential role of nitrate in accepting excess photochemically generated electrons, *Prochlorococcus* MIT0915 and *Prochlorococcus* MIT0917 cells were grown under nitrogen replete conditions, washed of residual nitrate, and then subjected to a nitrate pulse in the presence of light and temperature shocks. The maximum quantum yield of photochemistry at photosystem II (Fv/Fm) was used as a proxy for photochemical stress in the cells (Table 1). The control cultures exhibited an overall increase in Fv/Fm over the course of the experiment, while the cultures subjected to light and cold shocks had a lower Fv/Fm relative to each paired control. Most perturbations caused an overall decline in Fv/Fm over the 150 time course. These data indicate that each stressor elevated photochemical stress.

**Table 1.** Maximum quantum yield of photochemistry at photosystem II (Fv/Fm) for *Prochlorococcus* cells subjected to light and cold perturbations. Bolded values for Δ Fv/Fm indicate conditions where there was a decline in Fv/Fm over the 150 minute experiment.

| Experiment | Condition | $\mu\text{mol photons m}^{-2} \text{ s}^{-1}$ | Temp $^{\circ}\text{C}$ | Fv/Fm (0 min) | Fv/Fm (150 min) | $\Delta$ Fv/Fm |
| --- | --- | --- | --- | --- | --- | --- |
| *Prochlorococcus* MIT0915 |  |  |  |  |  |  |
| Light | Control | 22 | 24 | 0.417 (± 0.016) | 0.511 (± 0.015) | 0.094 |
| Light | Shock | 220 | 24 | 0.409 (± 0.017) | 0.272 (± 0.011) | **-0.137** |
| Cold | Control | 22 | 24 | 0.366 (± 0.009) | 0.486 (± 0.013) | 0.120 |
| Cold | Shock | 22 | 15 | 0.304 (± 0.010) | 0.226 (± 0.015) | **-0.078** |
| *Prochlorococcus* MIT0917 |  |  |  |  |  |  |
| Light | Control | 22 | 24 | 0.457 (± 0.004) | 0.539 (± 0.007) | 0.082 |
| Light | Shock | 220 | 24 | 0.463 (± 0.002) | 0.388 (± 0.005) | **-0.074** |
| Cold | Control | 22 | 24 | 0.427 (± 0.008) | 0.536 (± 0.009) | 0.108 |
| Cold | Shock | 22 | 15 | 0.385 (± 0.010) | 0.431 (± 0.002) | 0.046 |

Our hypothesis that nitrate reduction to nitrite could be enhanced as a response to photochemical stress was then evaluated by comparing nitrite production rates in the “shocked” cultures relative to the control cultures. At the original light and temperature conditions, all control cultures behaved as expected based on prior observations (Berube et al., 2023) — the MIT0915 strain did not produce nitrite following a nitrate pulse, while the MIT0917 strain did (Figure 2). The MIT0917 control cultures produced nitrite at rates of 1.04 × 10^−8^ (± 0.13 × 10^−8^) nmol nitrite cell^-1^ d^-1^ in the light shock experiment and 1.04 × 10^−8^ (± 0.17 × 10^−8^) nmol nitrite cell^-1^ d^-1^ in the cold shock experiment (Figure 2).

**Figure 2.**
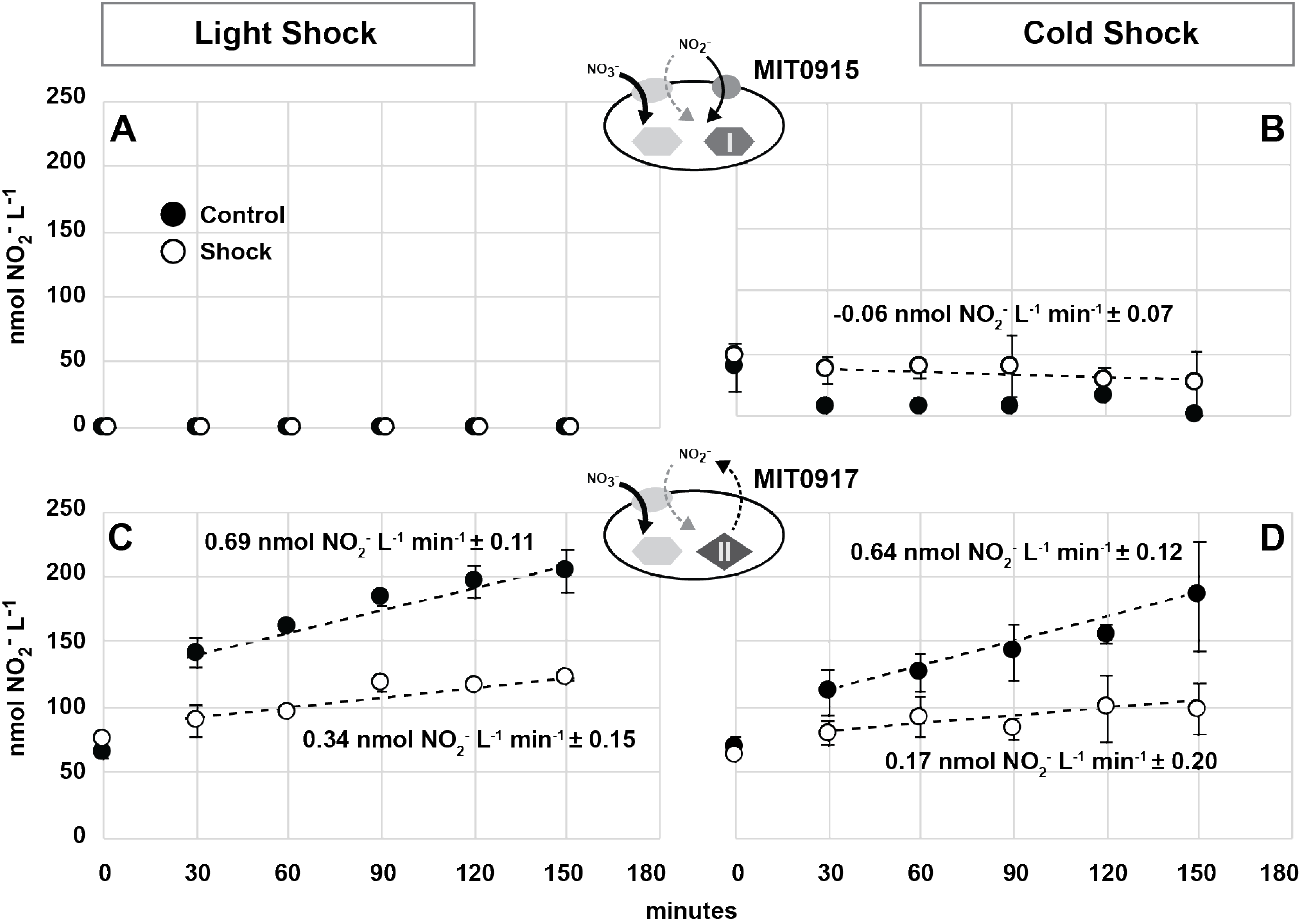
Nitrite production by *Prochlorococcus* during light and cold shocks. Dissolved nitrite concentrations were measured as a function of time in cultures of *Prochlorococcus* MIT0915 (A, B) and MIT0917 (C, D) following light (A, C) and temperature (B, D) perturbations. Cultures were shifted from a control photon flux of 22 μmol photons m^-2^ s^-1^ (A, C; closed circles) to a “shock” photon flux of 220 μmol photons m^-2^ s^-1^ (A, C; open circles) as well from a control temperature of 24°C (B, D; closed circles) to a “shock” temperature of 15°C (B, D; open circles). Error bars represent the standard deviation of 3 biological replicates. Rates are expressed as the mean ± one standard deviation calculated from each independent biological replicate. Where rates are not shown, it is because the majority of data points were below the nitrite detection limit.

The effect of both light and temperature perturbations on nitrite production rates demonstrate that increased photochemical electron pressure does not lead to elevated use of nitrate as an electron acceptor and concomitant release of nitrite as a waste product (Figure 2). Increasing the light intensity by over an order of magnitude decreased the cell-specific nitrite production rate of MIT0917 by 2.1 fold to 5.01 × 10^−9^ (± 2.04 × 10^−9^) nmol nitrite cell^-1^ d^-1^. Cold shock resulted in a more dramatic decline of nitrite production rates; at the lower temperature, the rate decreased by 3.8 fold to 2.76 × 10^−9^ (± 3.25 × 10^−9^) nmol nitrite cell^-1^ d^-1^. While we cannot rule out some degree of nitrite production in aiding the response of cells to stress — e.g., in the MIT0917 cells grown under light shock — we note that MIT0915 did not produce nitrite during either the light shock or cold shock experiments. Further, nitrite production in MIT0917 was relatively minimal during cold shock, consistent with the temperature dependence of nitrate uptake (Reay et al., 1999), rather than the use of nitrate as an escape valve for electrons to mitigate photochemical stress.

### Significance of enhanced nitrite production potential for nitrogen-limited *Prochlorococcus*

Overall, we observed a strong relationship between nitrite production following a nitrate pulse and nitrogen-limited physiology. This relationship was particularly apparent for the MIT0917 strain of *Prochlorococcus*, which lacks the FocA nitrite-specific transporter and encodes a divergent NirA nitrite reductase. Here, it is useful to compare contemporaneous experiments which used the same base artificial medium for nitrogen-limited (Figure 1) and nitrogen-replete (Figure 2) cultures. We observed that nitrite production rates following a nitrate pulse were more than an order of magnitude higher in nitrogen-limited cells of MIT0917 grown in chemostats (Figure 1) relative to nitrogen-replete cells in the control cultures used for the light and temperature shock experiments (Figure 2). When subjected to a pulse of nitrate, cells of MIT0917 that have been acclimated to N-limited growth exhibited 15-36 fold higher nitrite production rates than cells that were acclimated to N-replete conditions (Figure 3). Sequencing confirmed that the MIT0917 chemostats remained pure throughout the course of the experiment. While the MIT0915 chemostats were contaminated with *Aurantimonas*, our results suggest that the functional type represented by this strain may also release nitrite when challenged with a pulse of nitrate while in a nitrogen-limited state.

**Figure 3.**
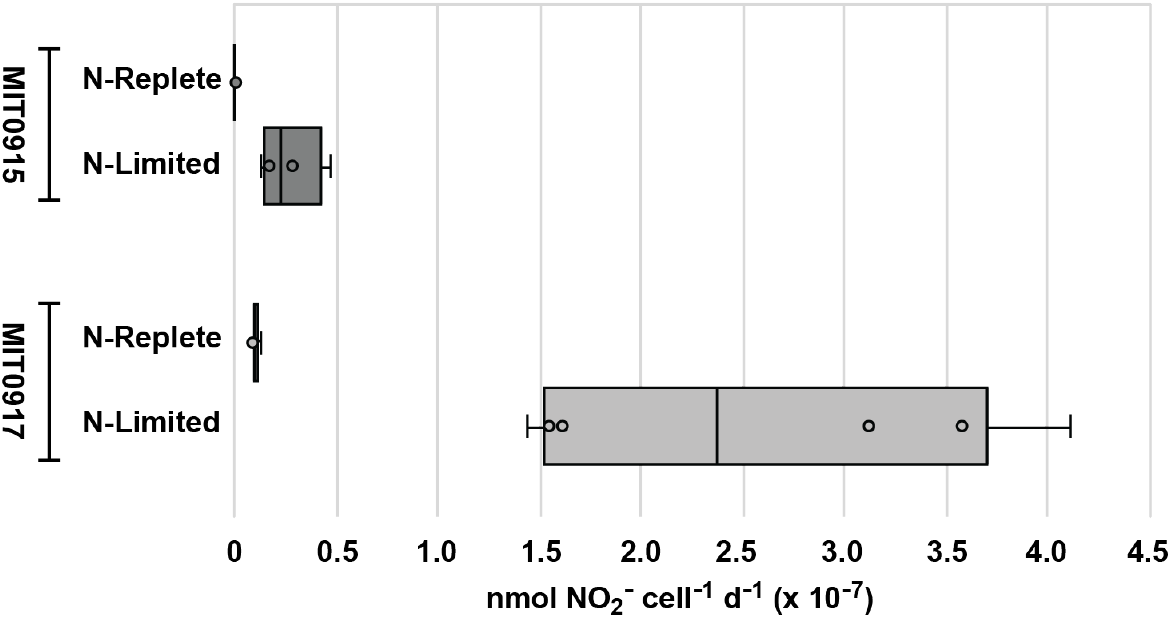
Comparison of nitrite production rates for nitrogen-replete and nitrogen-limited *Prochlorococcus* in response to a nitrate pulse. The rates for strains grown under nitrogen-replete conditions in artificial seawater are a compilation of rates for the control cultures in the light and temperature perturbation experiments (Figure 2). Rates for nitrogen-limited cultures were determined using sequential daily time points of nitrite concentration in duplicate chemostats following a nitrate pulse (Figure 1).

Perhaps surprisingly, we did not observe a relationship between nitrite production and stressors that are expected to increase photochemical electron pressure in *Prochlorococcus* cells (Figure 2). While this feature has been observed in larger size classes of phytoplankton (Glibert et al., 2013; Lomas & Glibert, 1999, 2000), LLI *Prochlorococcus* are known to be capable of tolerating photochemical stress (Malmstrom et al., 2010). In part, it is thought that the high number of genes encoding high-light inducible (Hli) or chlorophyll a/b-binding (CAB) proteins may facilitate the robust response of LLI *Prochlorococcus* to light shock (Coleman & Chisholm, 2007; Thompson, 2015). It has been hypothesized that internal waves or mixing events combined with changes in nitrate availability (i.e., seasonal upwellings in the Western Sargasso Sea) could contribute to nitrite release by phytoplankton due to an increase in photon flux density (Lomas & Lipschultz, 2006). Our data, however, suggest that the LLI *Prochlorococcus* in these systems may be more responsive to changes in nitrate availability rather than changes in photon flux.

In the context of cells in the wild, our results indicate that the magnitude and temporal variability of nitrite production by *Prochlorococcus* may be higher than expected. The magnitude of nitrite release appears to be especially high for cells starting from a nitrogen-limited state (Figure 3), and the timing of nitrite release by wild *Prochlorococcus* may indeed be dependent on the temporal variability nitrate pulses in marine ecosystems. It is conceivable that high nitrite production might primarily occur in cells living in nitrogen poor environments where there are intermittent pulses of nitrate from deeper waters — e.g., the North Pacific Subtropical Gyre (Johnson et al., 2010). LLI *Prochlorococcus* cells shifted to higher light intensities — e.g., due to internal waves that result in the shoaling of isopycnal layers — might not use nitrate as a photoprotective mechanism as observed for some other phytoplankton (Glibert et al., 2013; Lomas & Glibert, 1999, 2000). These observations add another dimension to our understanding of nitrogen cycling in oligotrophic, marine systems whereby perturbations in ambient nitrate concentrations may initiate a cascade of alternative N sources becoming available to other organisms due to the actions of abundant low-light adapted *Prochlorococcus*.

## Materials and Methods

### Strains and media

The cultures examined in this study included the MIT0915 and MIT0917 strains of *Prochlorococcus*, which both belong to the LLI clade of *Prochlorococcus* and are capable of using both nitrate and nitrite as their sole nitrogen source (Berube et al., 2023). All batch cultures were grown in AMP1-Mo-NO3 medium, which is a modified version of AMP1 artificial seawater medium (L. R. Moore et al., 2007) that was adjusted to contain 0.8 mmol L^-1^ sodium nitrate instead of 0.4 mmol L^-1^ ammonium sulfate and 100 nmol L^-1^ sodium molybdate instead of 0.3 nmol L^-1^. The higher molybdate concentration is needed to support the biosynthesis of the molybdopterin co-factor required for assimilatory nitrate reduction; 100 nmol L^-1^ is roughly the concentration of this trace metal in natural seawater (Horner et al., 2021; A. W. Morris, 1975). The medium was buffered using 5 mmol L^-1^ TAPS (pH=8.2) instead of HEPES (pH=7.5) and filter sterilized using a 0.2 μm polyethersulfone (PES) membrane filter instead of using steam sterilization in an autoclave to minimize heat induced hydrogen peroxide formation. In all experiments, 2.5 mmol L^-1^ of sodium bicarbonate was added as a carbon source. A nitrogen-free variant of AMP1-Mo-NO3 medium (AMP1-Mo) was also prepared by omitting all inorganic nitrogen sources and used for washing cells as described below. For the growth of *Prochlorococcus* in chemostats, the AMP1-Mo-NO3 medium was modified to have a 5:1 nitrogen:phosphorus ratio at final concentrations of 0.040 mmol L^-1^ sodium nitrate and 0.008 mmol L^-1^ sodium phosphate.

### Response of nitrate-limited cells to nitrate

The MIT0915 and MIT0917 strains of *Prochlorococcus* were grown under nitrate-limiting conditions in chemostats at dilution rates of 0.25 and 0.5 d^-1^ (Figure S1). Cell concentrations were routinely determined using flow cytometry, and undetectable levels of residual nitrate were confirmed using a continuous segmented flow autoanalyzer (see Supplementary Information). Cell-specific nitrite production rates for nitrate-limited *Prochlorococcus* MIT0915 and *Prochlorococcus* MIT0917 were determined following a nitrate pulse when the cultures were at a steady state at a dilution rate of 0.5 d^-1^. The nitrate pulse was initiated by aseptically adding 4 mL of 40 mmol L^-1^ sodium nitrate to each 200 mL chemostat culture. This yielded an initial concentration of 0.8 mmol L^-1^ of nitrate, with the nitrate concentration predicted to decline to below 0.05 mmol L^-1^ after 4 days at a dilution rate of 0.5 d^-1^ and assuming some consumption of the nitrate by the biomass. Samples for flow cytometry and nitrite analysis were collected immediately before the nitrate pulse and once per day following the nitrate pulse. The cell-specific nitrite production rate (NPR; nmol NO_2_^-^ cell^-1^ d^-1^) between sequential time points was determined using the following equation:

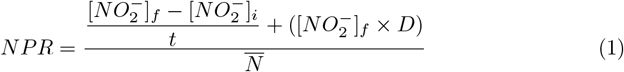

where 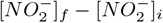 is the change in nitrite concentration between successive time points in units of nmol nitrite L^-1^, *t* is the time in days between the successive time points, 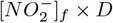 is the loss of nitrite from the chemostat at dilution rate, *D*, in units of nmol nitrite L^-1^ d^-1^, and 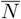 is the mean cell concentration for the two successive time points in cells L^-1^.

### Light and temperature shock experiments

MIT0915 and MIT0917 cells were grown in AMP1-Mo-NO3 at 20 μmol photons m^-2^ s^-1^ of blue light at 24°C until late exponential phase. Growth rates measured by fluorometry were 0.561 d^-1^ (± 0.073) for MIT0915 (N = 36) and 0.785 d^-1^ (± 0.030) for MIT0917 (N = 36). The cultures were harvested by centrifugation, washed of residual nitrate using AMP1-Mo, and resuspended to a final cell concentration of 1 × 10^−8^ cells mL^-1^ in nitrogen-free AMP1-Mo. For each experiment, triplicate biological cultures were prepared by pooling biomass from replicate batch cultures as described in the Supplementary Materials and Methods. The shock experiments were conducted in light- and temperature-controlled water baths, where the control conditions were maintained at 23–25°C and 21–23 μmol photons m^-2^ s^-1^ of blue light. In the light shock experiment, experimental cultures were exposed to 220 μmol photons m^-2^ s^-1^ of broad-spectrum light at 23-25°C. In the cold shock experiment, the cultures were exposed to 21–23 μmol photons m^-2^ s^-1^ of blue light at 14–16°C. Prior to spiking with nitrate, an initial set of samples was collected for flow cytometry, nitrite concentration determination, and fluorescence induction and relaxation (FIRe) fluorometry as described below. Nitrate was then added to each tube at a final concentration of 80 μmol L^-1^ sodium nitrate. Samples for nitrite concentration determination were collected every 30 min for 150 min and stored at −20°C. Endpoint flow cytometry and FIRe fluorometry samples were also collect at the final time point. Media-only control tubes were prepared using the same batch of nitrogen-free AMP1-Mo medium to facilitate assessment of background signal in nitrite concentration determination; the mean background signal was equivalent to 0.050 μmol L^-1^ nitrite (± 0.004) for the light shock experiment and 0.076 μmol L^-1^ nitrite (± 0.012) for the cold shock experiment.

## Acknowledgments

This work was supported by grants from the National Science Foundation (OCE-2048470 to P.M.B.), the Simons Foundation (Life Sciences Project Award ID 337262, S.W.C.; SCOPE Award ID 329108, S.W.C.), and the Robert and Ardis James Foundation to S.W.C. We thank the MIT BioMicro Center for assistance with DNA sequencing.

## Data Availability

Cell concentration, NO_2_^-^ concentration, fluorometry, and flow cytometry data for the light shock, cold shock, and continuous culture experiments are available under Project 833527 from the Biological and Chemical Oceanography Data Management. Sequencing data are available from the NCBI Sequence Read Archive under accessions SRX28796667, SRX28796668, SRX28813273, and SRX28813274.

## Author Contributions

**Paul M. Berube**: Conceptualization, Data curation, Formal analysis, Funding acquisition, Investigation, Methodology, Project administration, Supervision, Validation, Visualization, Writing – original draft, Writing – review and editing; **Trent LeMaster**: Data curation, Investigation, Methodology, Validation, Visualization, Writing – original draft, Writing – review and editing; **Sallie W. Chisholm**: Funding acquisition, Resources, Supervision, Writing – review and editing

## References

Bagby, S. C., & Chisholm, S. W. (2015). Response of Prochlorococcus to varying CO2:O2 ratios. The ISME Journal, 9 (10), 2232–2245. 10.1038/ismej.2015.36

Berube, P. M., Biller, S. J., Kent, A. G., Berta-Thompson, J. W., Roggensack, S. E., Roache-Johnson, K. H., Ackerman, M., Moore, L. R., Meisel, J. D., Sher, D., Thompson, L. R., Campbell, L., Martiny, A. C., & Chisholm, S. W. (2015). Physiology and evolution of nitrate acquisition in Prochlorococcus. The ISME Journal, 9 (5), 1195–1207. 10.1038/ismej.2014.211

Berube, P. M., O’Keefe, T. J., Rasmussen, A., LeMaster, T., & Chisholm, S. W. (2023). Production and cross-feeding of nitrite within Prochlorococcus populations. mBio, 0 (0), e01236–23. 10.1128/mbio.01236-23

Berube, P. M., Rasmussen, A., Braakman, R., Stepanauskas, R., & Chisholm, S. W. (2019). Emergence of trait variability through the lens of nitrogen assimilation in Prochlorococcus (P. G. Falkowski, I. T. Baldwin, & J. Morris, Eds.). eLife, 8, e41043. 10.7554/eLife.41043

Biller, S. J., Berube, P. M., Lindell, D., & Chisholm, S. W. (2015). Prochlorococcus: The structure and function of collective diversity. Nature Reviews Microbiology, 13 (1), 13–27. 10.1038/nrmicro3378

Braakman, R., Follows, M. J., & Chisholm, S. W. (2017). Metabolic evolution and the self-organization of ecosystems. Proceedings of the National Academy of Sciences, 114 (15). 10.1073/pnas.1619573114

Coleman, M. L., & Chisholm, S. W. (2007). Code and context: Prochlorococcus as a model for cross-scale biology. Trends in Microbiology, 15 (9), 398–407. 10.1016/j.tim.2007.07.001

Coleman, M. L., & Chisholm, S. W. (2010). Ecosystem-specific selection pressures revealed through comparative population genomics. Proceedings of the National Academy of Sciences, 107 (43), 18634–18639. 10.1073/pnas.1009480107

Droop, M. R. (1968). Vitamin B12 and Marine Ecology. IV. The Kinetics of Uptake, Growth and Inhibition in Monochrysis lutheri. Journal of the Marine Biological Association of the United Kingdom, 48 (3), 689–733. 10.1017/S0025315400019238

Droop, M. R. (1973). Some Thoughts on Nutrient Limitation in Algae. Journal of Phycology, 9 (3), 264–272. 10.1111/j.1529-8817.1973.tb04092.x

Droop, M. R. (1974). The nutrient status of algal cells in continuous culture. Journal of the Marine Biological Association of the United Kingdom, 54 (4), 825–855. 10.1017/S002531540005760X

Flombaum, P., Gallegos, J. L., Gordillo, R. A., Rincón, J., Zabala, L. L., Jiao, N., Karl, D. M., Li, W. K. W., Lomas, M. W., Veneziano, D., Vera, C. S., Vrugt, J. A., & Martiny, A. C. (2013). Present and future global distributions of the marine Cyanobacteria Prochlorococcus and Synechococcus. Proceedings of the National Academy of Sciences, 110 (24), 9824–9829. 10.1073/pnas.1307701110

Garcia, C. A., Hagstrom, G. I., Larkin, A. A., Ustick, L. J., Levin, S. A., Lomas, M. W., & Martiny, A. C. (2020). Linking regional shifts in microbial genome adaptation with surface ocean biogeochemistry. Philosophical Transactions of the Royal Society B: Biological Sciences, 375 (1798), 20190254. 10.1098/rstb.2019.0254

Glibert, P. M., Kana, T. M., & Brown, K. (2013). From limitation to excess: The consequences of substrate excess and stoichiometry for phytoplankton physiology, trophodynamics and biogeochemistry, and the implications for modeling. Journal of Marine Systems, 125, 14–28. 10.1016/j.jmarsys.2012.10.004

Horner, T. J., Little, S. H., Conway, T. M., Farmer, J. R., Hertzberg, J. E., Janssen, D. J., Lough, A. J. M., McKay, J. L., Tessin, A., Galer, S. J. G., Jaccard, S. L., Lacan, F., Paytan, A., Wuttig, K., & Members, G.-P. B. P. W. G. (2021). Bioactive Trace Metals and Their Isotopes as Paleoproductivity Proxies: An Assessment Using GEOTRACES-Era Data. Global Biogeochemical Cycles, 35 (11), e2020GB006814. 10.1029/2020GB006814

Johnson, K. S., Riser, S. C., & Karl, D. M. (2010). Nitrate supply from deep to near-surface waters of the North Pacific subtropical gyre. Nature, 465 (7301), 1062–1065. 10.1038/nature09170

Kamennaya, N. A., Chernihovsky, M., & Post, A. F. (2008). The cyanate utilization capacity of marine unicellular Cyanobacteria. Limnology and Oceanography, 53 (6), 2485–2494. 10.4319/lo.2008.53.6.2485

Kamennaya, N. A., & Post, A. F. (2011). Characterization of Cyanate Metabolism in Marine Synechococcus and Prochlorococcus spp. Applied and Environmental Microbiology, 77 (1), 291–301. 10.1128/AEM.01272-10

Lomas, M. W., & Glibert, P. M. (1999). Temperature regulation of nitrate uptake: A novel hypothesis about nitrate uptake and reduction in cool-water diatoms. Limnology and Oceanography, 44 (3), 556–572. 10.4319/lo.1999.44.3.0556

Lomas, M. W., & Glibert, P. M. (2000). Comparisons of Nitrate Uptake, Storage, and Reduction in Marine Diatoms and Flagellates. Journal of Phycology, 36 (5), 903–913. 10.1046/j.1529-8817.2000.99029.x

Lomas, M. W., & Lipschultz, F. (2006). Forming the primary nitrite maximum: Nitrifiers or phytoplankton? Limnology and Oceanography, 51 (5), 2453–2467. 10.4319/lo.2006.51.5.2453

Malmstrom, R. R., Coe, A., Kettler, G. C., Martiny, A. C., Frias-Lopez, J., Zinser, E. R., & Chisholm, S. W. (2010). Temporal dynamics of Prochlorococcus ecotypes in the Atlantic and Pacific oceans. The ISME Journal, 4 (10), 1252–1264. 10.1038/ismej.2010.60

Martiny, A. C., Coleman, M. L., & Chisholm, S. W. (2006). Phosphate acquisition genes in Prochlorococcus ecotypes: Evidence for genome-wide adaptation. Proceedings of the National Academy of Sciences, 103 (33), 12552–12557. 10.1073/pnas.0601301103

Moore, C. M., Mills, M. M., Arrigo, K. R., Berman-Frank, I., Bopp, L., Boyd, P. W., Galbraith, E. D., Geider, R. J., Guieu, C., Jaccard, S. L., Jickells, T. D., La Roche, J., Lenton, T. M., Mahowald, N. M., Marañón, E., Marinov, I., Moore, J. K., Nakatsuka, T., Oschlies, A., … Ulloa, O. (2013). Processes and patterns of oceanic nutrient limitation. Nature Geoscience, 6 (9), 701–710. 10.1038/ngeo1765

Moore, L. R., Coe, A., Zinser, E. R., Saito, M. A., Sullivan, M. B., Lindell, D., Frois-Moniz, K., Waterbury, J., & Chisholm, S. W. (2007). Culturing the marine cyanobacterium Prochlorococcus. Limnology and Oceanography: Methods, 5 (10), 353–362. 10.4319/lom.2007.5.353

Moore, L. R., Post, A. F., Rocap, G., & Chisholm, S. W. (2002). Utilization of different nitrogen sources by the marine cyanobacteria Prochlorococcus and Synechococcus. Limnology and Oceanography, 47 (4), 989–996. 10.4319/lo.2002.47.4.0989

Morris, A. W. (1975). Dissolved molybdenum and vanadium in the northeast Atlantic Ocean. Deep Sea Research and Oceanographic Abstracts, 22 (1), 49–54. 10.1016/0011-7471(75)90018-2

Morris, J. J., Johnson, Z. I., Szul, M. J., Keller, M., & Zinser, E. R. (2011). Dependence of the Cyanobacterium Prochlorococcus on Hydrogen Peroxide Scavenging Microbes for Growth at the Ocean’s Surface. PLOS ONE, 6 (2), e16805. 10.1371/journal.pone.0016805

Morris, J. J., Kirkegaard, R., Szul, M. J., Johnson, Z. I., & Zinser, E. R. (2008). Facilitation of Robust Growth of Prochlorococcus Colonies and Dilute Liquid Cultures by “Helper” Heterotrophic Bacteria. Applied and Environmental Microbiology, 74 (14), 4530–4534. 10.1128/AEM.02479-07

Morris, J. J., Lenski, R. E., & Zinser, E. R. (2012). The Black Queen Hypothesis: Evolution of Dependencies through Adaptive Gene Loss. mBio, 3 (2), 10.1128/mbio.00036–12. 10.1128/mbio.00036-12

Reay, D. S., Nedwell, D. B., Priddle, J., & Ellis-Evans, J. C. (1999). Temperature Dependence of Inorganic Nitrogen Uptake: Reduced Affinity for Nitrate at Suboptimal Temperatures in Both Algae and Bacteria. Applied and Environmental Microbiology, 65 (6), 2577–2584. 10.1128/AEM.65.6.2577-2584.1999

Shimakawa, G., & Miyake, C. (2018). Oxidation of P700 Ensures Robust Photosynthesis. Frontiers in Plant Science, 9. 10.3389/fpls.2018.01617

Thompson, J. W. (2015). Prochlorococcus : Life in light [Thesis]. Massachusetts Institute of Technology. Retrieved May 7, 2025, from https://dspace.mit.edu/handle/1721.1/99797 xAccepted: 2015-11-09T19:49:31Z.

Ustick, L. J., Larkin, A. A., Garcia, C. A., Garcia, N. S., Brock, M. L., Lee, J. A., Wiseman, N. A., Moore, J. K., & Martiny, A. C. (2021). Metagenomic analysis reveals global-scale patterns of ocean nutrient limitation. Science, 372 (6539), 287–291. 10.1126/science.abe6301

Yelton, A. P., Acinas, S. G., Sunagawa, S., Bork, P., Pedrós-Alió, C., & Chisholm, S. W. (2016). Global genetic capacity for mixotrophy in marine picocyanobacteria. The ISME Journal, 10 (12), 2946–2957. 10.1038/ismej.2016.64

Zubkov, M. V., Tarran, G. A., & Fuchs, B. M. (2004). Depth related amino acid uptake by Prochlorococcus cyanobacteria in the Southern Atlantic tropical gyre. FEMS Microbiology Ecology, 50 (3), 153–161. 10.1016/j.femsec.2004.06.009

