## Supplementary Appendix for "Priming the pump: Enhanced nitrite release in response to a nitrate pulse by nitrogen-limited *Prochlorococcus*"

### Supplementary Materials and Methods

***Prochlorococcus* strain maintenance.** The MIT0915 and MIT0917 strains of *Prochlorococcus* were routinely monitored for purity by using a panel of purity broths (Berube et al., 2015) as well as flow cytometry of SYBR green stained cultures to identify the potential presence of heterotrophic (non-chlorophyll containing) bacteria. Sanger sequencing of the 16S-23S intergenic spacer sequence (ITS) was used to confirm strain identity before experiments. Additionally, short-read sequencing was used, as detailed below, to assess the purity of continuous cultures at the conclusion of the experiment given the difficulty of aseptically sampling the chemostats from the outflow pump. All cultures were grown for at least 10 generations to ensure acclimation to growth conditions as determined by consistent growth rates assessed by relative fluorescence measured using a Turner TD-700 Fluorometer.

**Chemostat design and operation.** *Prochlorococcus* MIT0915 and *Prochlorococcus* MIT0917 were grown in chemostats with nitrate as the limiting nutrient in AMP1-Mo-NO3 artificial medium adjusted to have a 5:1 nitrogen to phosphorus ratio (0.040 mmol $L^{-1}$ sodium nitrate and 0.008 mmol $L^{-1}$ sodium phosphate). Chemostat cultures were grown at 24°C and 26 µmol photons $m^{-2}$ $s^{-1}$ (± 2) of blue light in a custom designed continuous cultivation system for picocyanobacteria (Figure S1). The chemostat vessels — constructed from 35 mm x 300 mm glass hybridization bottles — were placed in a glass walled aquarium with temperature controlled by an aquarium heater and light filtered through #69 brilliant blue gel film from LED light banks. Freshly prepared medium was sterile filtered through a 0.2 µm polyethersulfone (PES) sterivex filter directly into sterile 5 L glass bottles. Two dedicated high precision peristaltic pumps (Golander, Norcross, GA) were used for pumping media into the chemostats (inflow pump) and pumping sample out of the chemostats (outflow pump), respectively. The volume of the chemostats was controlled through positive pressure coupled with an overflow tube placed at an empirically determined level to maintain a volume of 200 mL. Mixing, positive pressure, and aeration was provided by bubbling humidified air through a 0.2 µm PTFE venting filter. Before reaching the chemostat vessels, the air was sequentially sparged through 18 MΩ·cm water to humidify, 1 N sulfuric acid to trap gaseous ammonia, and again through 18 MΩ·cm water to trap any residual acid. The cultures were grown as duplicates at dilution rates of 0.25 $d^{-1}$ for days 1-106 and 0.5 $d^{-1}$ for days 107-162 in order to confirm expected changes in cell numbers in response to imposed growth rates. Cell concentrations were monitored three times per week using flow cytometry as described below. Low background samples for nitrite and nitrate analysis were collected by pumping culture into an acid-washed vessel, centrifuged in 35 mL acid-washed oakridge tubes at 12,500 RPM in a JA-25.50 rotor for 25 min to pellet biomass, with the supernatant retained and stored at -20°C for no longer than 30 days before nitrite and/or nitrate analysis.

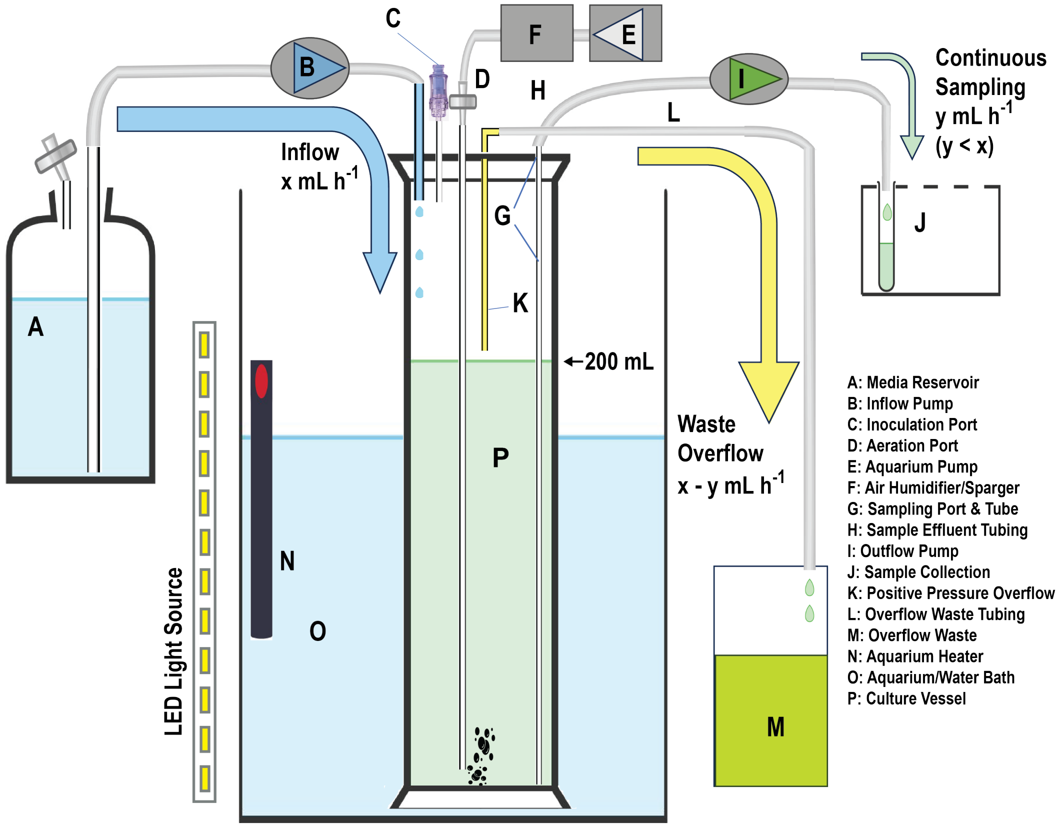


**Figure 1. Design schematic for chemostats used to grow *Prochlorococcus* under nitrate-limitation.** A low-cost light and temperature controlled continuous cultivation system was developed for nutrient-limited growth of *Prochlorococcus*. This schematic was made in part using Chemix.

**Assessment of continuous culture purity.** Short-read sequencing was used to assess the purity of each chemostat at the conclusion of the continuous culturing experiment. The chemostat cultures were decanted into a sterile 250 mL polycarbonate centrifuge bottle and then centrifuged at 12,000 RPM in a JA-14 Rotor for 25 min. The pellet was resuspended in 100 µL of AMP1-Mo-NO3 and then stored at -80°C. DNA was prepared from the frozen biomass using phenol/chloroform extraction (Wilson, 2001). Approximately 4-5 million 150+150 nt paired end reads were generated for each sample using the Illumina NovaSeq platform by the MIT BioMicro Center. The bbmap package (version 39) was used for adapter trimming, quality filtering, and mapping of reads to the *Prochlorococcus* MIT0915 and *Prochlorococcus* MIT0917 genomes. Unmapped reads derived from the MIT0915 chemostat samples — associated with heterotrophic bacterial contaminants — were assembled using SPAdes, binned using metabat, annotated with prokka, and taxonomically classified using gtdbtk. Read mapping using bbmap was repeated using the assembled heterotroph genomes as additional reference sequences to assess the proportion of reads mapping to these contaminants. The MIT0917 culture remained pure throughout the course of chemostat operation given that >98% of reads mapped to the MIT0917 genome. The MIT0915 culture, however, contained *Aurantimonas* (38-41% of reads), *Mycobacterium* (3-10% of reads), and *Nocardioides* (3-4% of reads), with 43% and 52% of reads mapping to *Prochlorococcus* MIT0915 in chemostats A and B, respectively.

**Biomass generation for light and temperature perturbation experiments.** *Prochlorococcus* MIT0915 and *Prochlorococcus* MIT0917 cells were grown in sterile borosilicate glass culture tubes in 35 mL of AMP1-Mo-NO3 at 20 µmol photons $m^{-2}$ $s^{-1}$ of blue light at 24°C. Biomass was generated by growing 18 replicates of 35 mL cultures for each strain and then pooling 3 sets of 6 tubes to serve as 3 biological replicates. These replicate pooled cultures were centrifuged at 10,000 RPM for 20 minutes at 22°C using a JA-14 rotor. Cell pellets were resuspended and washed in 100 mL of nitrogen-free AMP1-Mo medium to remove residual nitrate, centrifuged again, and then resuspended in 50 mL of nitrogen-free AMP1-Mo. Cell concentrations for the resuspended biomass samples were then determined by flow cytometry. Prior to the initiation of the light and temperature perturbations, each biological replicate was diluted to 1 x $10^{-8}$ cells $mL^{-1}$ in a total of 50 mL of nitrogen-free AMP1-Mo supplemented with 2.5 mM sodium bicarbonate in duplicate — one to serve as a control and one to be subjected to the perturbation. For the temperature perturbation experiment, the nitrogen-free AMP1-Mo medium was pre-chilled to 15°C to ensure a rapid shift to the temperature shock condition.

**Flow cytometry.** Cell concentrations were determined by flow cytometry using a Guava easyCyte 12HT Flow Cytometer (MilliporeSigma, Burlington, MA, USA). Samples were either run as live cultures after dilution into fresh medium that was filtered using a 0.2 µm PES syringe filter or after fixation with 0.125% glutaraldehyde and storage at -80°C following flash-freezing in liquid nitrogen. *Prochlorococcus* cells were identified by forward scatter and red fluorescence excited by a 488 nm laser. All cultures were diluted to 50 to 200 cells $\mu L^{-1}$ and data were collected for up to 6 minutes at a flow rate of 0.024 $\mu L\ s^{-1}$. Guava easyCheck beads (MilliporeSigma, Burlington, MA, USA) were run daily to verify that the instrument was functioning within normal operating parameters and meeting predefined tolerances. For each run, a diluted sample of Guava easyCheck beads was run using the same parameters as the cultures to serve as a standard for the normalization of forward scatter and fluorescence data. Flow cytometry data were exported as FSC 3.0 formatted files and analyzed using FlowJo (BD Biosciences, Ashland, OR, USA).

**Nitrite and nitrate measurements.** Nitrite was analyzed using an AA3 HR Autoanalyzer (Seal Analytical, Milwaukee, WI, USA). In this continuous segmented flow analyzer, nitrite is reacted with sulfanilamide and N-(1-naphthyl)ethylenediamine (NED) to produce a pink diazo dye (Wood 1967; Hansen 1999). For determination of total nitrate and nitrite, any nitrate in the sample was first reduced to nitrite using a copper-cadmium coil. Absorbance was measured in a 1 cm flow cell using a 550 nm bandpass filter (SEAL Analytical method number G-384-08 for a multitest MT19B manifold). For each analysis, sodium nitrite and sodium nitrate standards were freshly prepared and serially diluted in nitrogen-free AMP1 medium to create a calibration curve at the following concentrations: 1.25, 0.5, 0.25, 0.08, 0.04, 0.03, and 0.015 µmol $L^{-1}$. A linear calibration curve was applied to the data and the instrument's response factor, represented by the slope of the calibration curve, remained consistent and reproducible throughout the nitrite analysis period ($r^2$ >0.99 in all runs). For samples in nitrogen-free AMP1 medium as a background matrix, the limit of detection (LOD) for nitrite was 0.010 µmol $L^{-1}$ and the limit of blank was 0.003 µmol $L^{-1}$. All samples were analyzed within one month of storage at -20°C.

**Fluorescence induction and relaxation**. The efficiency for energy conversion at photosystem II (Fv/Fm) for cultures subjected to light or cold shocks was measured using a FIRe Fluorometer System (Satlantic, Halifax, NS, Canada). A sample of each culture was diluted 10 fold in nitrogen-free AMP1 medium, incubated in the dark for 10 minutes in order to allow photosystems to open, and then transferred to a cylindrical quartz cuvette. A total of 10 profiles were collected for each sample. Fluorescence data were processed using fireworx 0.9.1.

### References

Berube, P. M., Biller, S. J., Kent, A. G., Berta-Thompson, J. W., Roggensack, S. E., Roache-Johnson, K. H., Ackerman, M., Moore, L. R., Meisel, J. D., Sher, D., Thompson, L. R., Campbell, L., Martiny, A. C., & Chisholm, S. W. (2015). Physiology and evolution of nitrate acquisition in *Prochlorococcus*. *The ISME Journal*, *9*(5), 1195–1207. https://doi.org/10.1038/ismej.2014.211

Wilson, K. (2001). Preparation of Genomic DNA from Bacteria. *Current Protocols in Molecular Biology*, *56*(1), 2.4.1–2.4.5. https://doi.org/10.1002/0471142727.mb0204s56
